# Pragmatic spatial sampling for wearable MEG arrays

**DOI:** 10.1101/2019.12.29.890426

**Authors:** Tim M Tierney, Stephanie Mellor, George C O’Neill, Niall Holmes, Elena Boto, Gillian Roberts, Ryan M Hill, James Leggett, Richard Bowtell, Matthew J Brookes, Gareth R Barnes

## Abstract

Several new technologies have recently emerged promising new MEG systems in which the sensors can be placed close to the scalp. One such technology, Optically Pumped Magnetometry MEG (OP-MEG) allows for a scalp mounted flexible system that provides field measurements within mm of the scalp surface. A question that arises in developing on-scalp systems, such as OP-MEG scanners, is: how many sensors are necessary to achieve adequate performance/spatial discrimination? There are many factors to consider in answering this question such as the signal to noise ratio (SNR), the locations and depths of the sources of interest, the density of spatial sampling, sensor gain errors (due to interference, subject movement, cross-talk, etc.) and, of course, the desired spatial discrimination. In this paper, we provide simulations which show the impact these factors have on designing sensor arrays for wearable MEG. While OP-MEG has the potential to provide high information content at dense spatial samplings, we find that adequate spatial discrimination of sources (<1cm) can be achieved with relatively few sensors (<100) at coarse spatial samplings (~30mm) at high SNR. Comparable discrimination for traditional cryogenic systems require far more channels by these same metrics. Finally we show that sensor gain errors have the greatest impact on discrimination between deep sources at high SNR.

## 1. Introduction

Magnetoencephalography (MEG) has become a vital tool for studying human brain function in both basic and clinical settings (Bagic, Bowyer, Kirsch, Funke, & Burgess, 2017; Baillet, 2017; Matti S Hämäläinen, Hari, Ilmoniemi, Knuutila, & Lounasmaa, 1993; Rampp et al., 2019). Simulation studies show that MEG system performance continues to improve with an increase in channel count as long as channel noise is uncorrelated (Jiri Vrba & Robinson, 2002). There is however a balance between system cost and channel redundancy. In the last 20 years the design of whole-head Superconducting Quantum Interference Device (SQUID) based systems stabilized to around 300 channels. This figure was arrived at after a great deal of early work to quantify the problem at hand (Ahonen et al., 1993; Kemppainen & Ilmoniemi, 1989; J. Vrba, Robinson, & McCubbin, 2004; Jiri Vrba & Robinson, 2002). Many of these design considerations assume perfect knowledge of the underlying current distribution, but there is also a complex interplay between the inversion assumptions, forward model, sampling density, and the geometry of the source space (Ahonen et al., 1993; Barnes, Hillebrand, Fawcett, & Singh, 2004; Brookes et al., 2010; Hauk, Stenroos, & Treder, 2019; Hedrich, Pellegrino, Kobayashi, Lina, & Grova, 2017; A. Hillebrand & Barnes, 2002; Iivanainen, Stenroos, & Parkkonen, 2017; Sekihara, Sahani, & Nagarajan, 2008).

These issues have come back into focus with emergence of new MEG system technologies. These include the advent of high critical temperature SQUIDs (high T_c_- SQUIDS), nitrogen vacancy magnetometers and optically pumped magnetometers (OPMs) (Allred, Lyman, Kornack, & Romalis, 2002; Faley et al., 2013; Hingant et al., 2014; Osborne, Orton, Alem, & Shah, 2018; Schneiderman, 2014; Tierney et al., 2019). These new sensors allow for the construction of wearable arrays that offer the flexibility to image human brain function during subject movement (Boto et al., 2018; Roberts et al., 2019; Tierney et al., 2018), with the crucial addition of field nulling coils (Holmes et al., 2018, 2019; Iivanainen, Zetter, Grön, Hakkarainen, & Parkkonen, 2019).

These sensors are also typically placed much closer to the scalp (a few mm) than traditional MEG sensors, resulting in a 3-5 fold increase in signal (Boto, Meyer, et al., 2016; Iivanainen et al., 2017). In principle, this should offer the capability of resolving neuronal activity with a higher spatial discrimination as smaller spatial wavelengths can potentially be sampled (Ahonen et al., 1993; Boto et al., 2019). This motivates sampling more densely to take advantage of this extra information, as has recently been suggested in a study of visual gamma using optically pumped magnetometers (Iivanainen, Zetter, & Parkkonen, 2019) and in a theoretical paper on spatial sampling (Iivanainen, Mäkinen, Zetter, et al., 2019) which suggests that on-scalp systems would benefit from 3 times as many sensors as off-scalp systems to maximise spatial discrimination.

Here we argue that for a wide variety of neuroscience applications maximising information content or spatial discrimination, although desirable, is not necessary. For instance, in paediatric epilepsy surgery, a crucial clinical application of MEG (De Tiège et al., 2012; Doss, Zhang, Risse, & Dickens, 2009; Englot et al., 2015; Rosenow & Lüders, 2001; Schneider et al., 2013; Sutherling, Mamelak, Thyerlei, & Maleeva, 2008) a large percentage of brain volume (~5-10%, equivalent to a 2-3cm radius sphere) may be resected (Centeno et al., 2017). The clinical question does not require the spatial discrimination between sources at a millimetre scale. In the case of basic neuroscience applications that average results across subjects have their spatial discrimination limited by the functional and anatomical variability that exists between subjects (Aquino et al., 2019; Devlin & Poldrack, 2007; Geyer & Turner, 2013; Sabuncu et al., 2010). Furthermore, in studies which investigate electrophysiological functional connectomes, the spatial discrimination required may be on the scale of 1-2 centimetres, as a single time course representing an entire atlas-defined parcel is often utilised (O’Neill et al., 2018). Rather than strive to maximize spatial discrimination or information content, an alternative question is “given a desired discrimination what is the sampling density/ sensor number I require?” We approach the question of discrimination in a statistical framework by asking “at which sensor density one is able to confidently (p<0.05) distinguish between two competing source models” (Meyer et al., 2017a; R.N. Henson, J. Mattout, C. Phillips, & K.J. Friston, 2009; Troebinger, López, Lutti, Bestmann, & Barnes, 2014). By adopting this approach we can design MEG arrays around a given scientific question (or spatial discrimination) and therefore minimize channel count.

Unlike SQUID based systems, where the feedback electronics maintains a constant linear relationship to applied flux optically pumped magnetometers are particularly vulnerable to external factors which give rise to a change in their gain. Perhaps the most pernicious of these errors is due to the inherent nonlinear response of the sensor (Tierney et al., 2019). When operating in the spin exchange relaxation Free (SERF) regime these sensors are only linear within a few nT of zero field (Boto et al., 2018). These gain errors can be caused by subject movement through a non-zero background field or by the change in the ambient magnetic field over time (Holmes et al., 2019; Iivanainen, Zetter, Grön, et al., 2019). These issues would typically require active shielding in order to be minimized. Another issue is how the sensor interact with each other in multichannel systems. On-board coils that produce magnetic fields for field zeroing or amplitude modulation (Cohen-Tannoudji, Dupont-Roc, Haroche, & Laloë, 1970; Osborne et al., 2018) may change the gain of a nearby sensor. This problem is static (if the sensors do not move relative to one another) and deterministic but assumes that a suitable calibration procedure is in place to correct for these issues.

The paper proceeds as follows. First we outline the model comparison framework and the competing models used to define spatial discrimination. We examine these discrimination estimates for different sensor numbers under different SNR conditions for deep and shallow dipolar sources using on-scalp sensors. We use the same methods to investigate optimal cryogenic sensors layouts and show, in accord with previous studies (Boto, Bowtell, et al., 2016), that comparable discrimination for on-scalp sensors is achievable with fewer sensors. Finally we examine the impact of sensor gain error (Boto et al., 2018; Iivanainen, Zetter, Grön, et al., 2019; Tierney et al., 2019) on discrimination. The conclusion is that for typical cognitive neuroscience or clinical requirements of 1cm discrimination, between 50-100 on-scalp sensors are required and that sensor gain errors have the greatest impact when the task is to distinguish between proximal deep sources.

## 2. Methods

### 2.1 Model comparison and spatial discrimination

Here we define a metric of spatial discrimination as the Euclidian distance at which we can confidently (p<0.05) distinguish between the magnetic field patterns generated by two sources on the cortical mesh. We use model comparison to compare data from these source models in terms of their variational free energy (Friston, Mattout, Trujillo-Barreto, Ashburner, & Penny, 2007). The method has 4 steps that are described graphically in Figure 1: 1) A source location is used to generate a single dataset. 2) Different generative models of the same data are proposed. The “base model” includes a source and all vertices within 20 mm of the source (top-left panel figure 1). All other models do not include the source (or the lead-field elements mapping sensors to the simulated location) and any potential sources (or lead-field mappings) within a specified radius. This radius (excluding the lead-field mappings from sensors to the true source and its neighbours) is gradually increased in 1mm steps (top panels figure 1). 3) Each candidate model is inverted (using the limited source space) to explain the source-generated dataset and a Free energy (or log model evidence) value is obtained. Importantly, only the base-model includes the generating source. 4) As the data remain the same we can compare the models used to describe the data probabilistically. The free energy difference between any two models being how likely (on log scale) one model is over the other. The ring model that is 20 times (or log difference of 3) less likely than the base model defines the spatial discrimination of the array. Similar approaches have been used elsewhere to define the discriminability of different hippocampal (Meyer et al., 2017a) and cortical mesh models. A more thorough introduction to the use of free energy in source inversion is given elsewhere (Lopez, Litvak, Espinosa, Friston, & Barnes, 2014).

**Figure 1.**
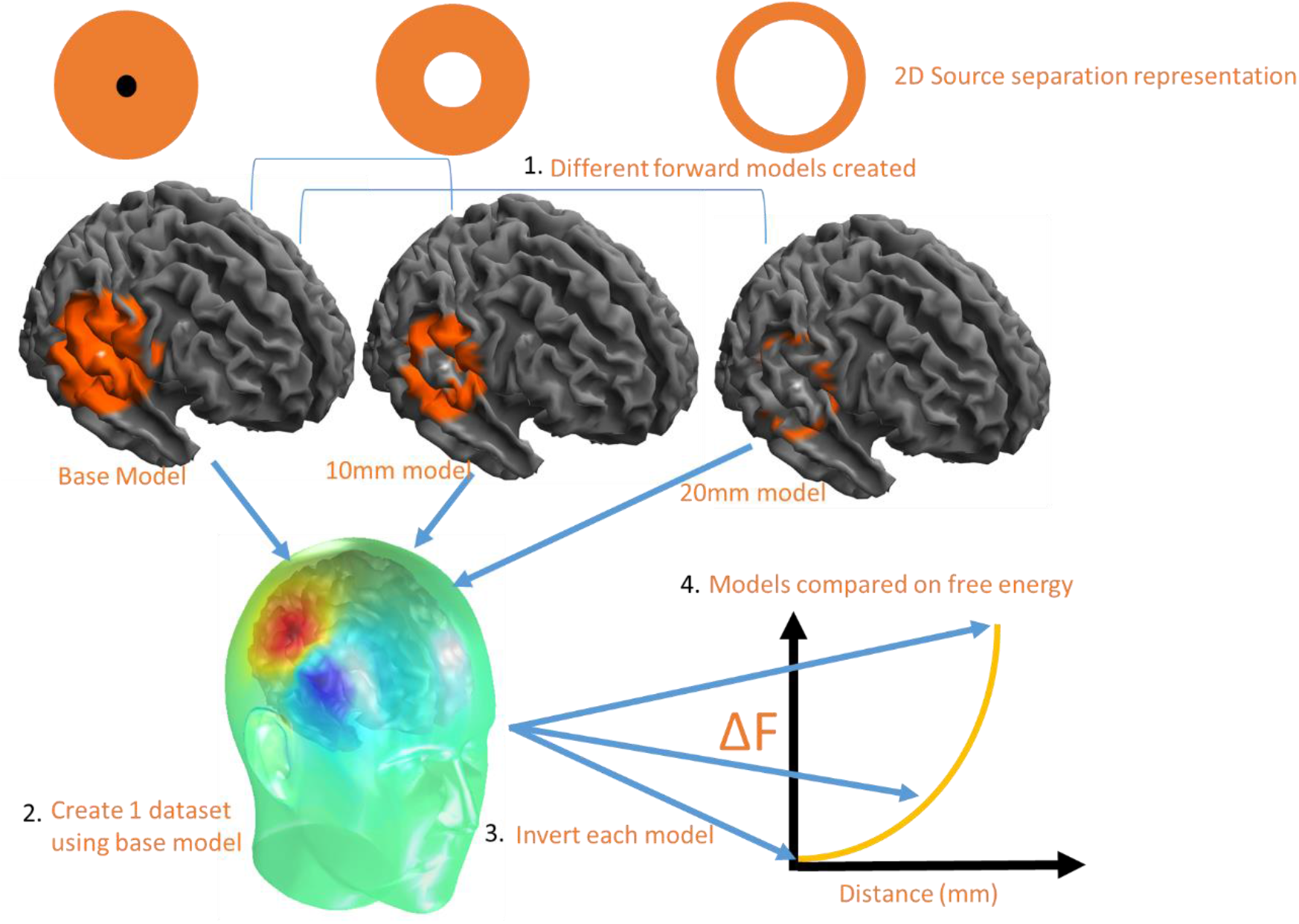
Model comparison and spatial discrimination. Different forward models are created that map to brain regions that are progressively more distant to the true source of the brain activity (signified in orange). A 2D representation is provided with the source in black and the candidate models in orange (the contents of the lead field matrix). We then reconstruct the simulated data onto all cortical models. Each fit has an associated model evidence or free energy value. We look for the model (ring radius from the true source) with a log free energy of 3 greater than the base model, indicating that we can confidently (p < 0.05) distinguish between the two.

### 2.2 Array Design

We use a point packing algorithm to position sensors on the scalp surface at increasing densities (e.g. 20, 15, 10 mm separation). There are 5 steps to the algorithm and the approach is depicted graphically in Figure 2. 1) Initially the bounding box of the surface is subdivided into squares of equal area with an edge length equal to the desired sampling density. 2) The corners of the squares are then projected onto the surface to initialise the algorithm at approximately the correct sampling density. 3) In the optimisation stage, at each loop iteration, a sensor is chosen at random and moved to a neighbouring vertex if this brings the sampling density closer to the target sampling density. 4) If there are any vertices on the surface that are farther from the nearest sensor than the target spacing, a sensor is added to that vertex. 5) This process is repeated for a large (~10,000) number of iterations.

**Figure 2.**
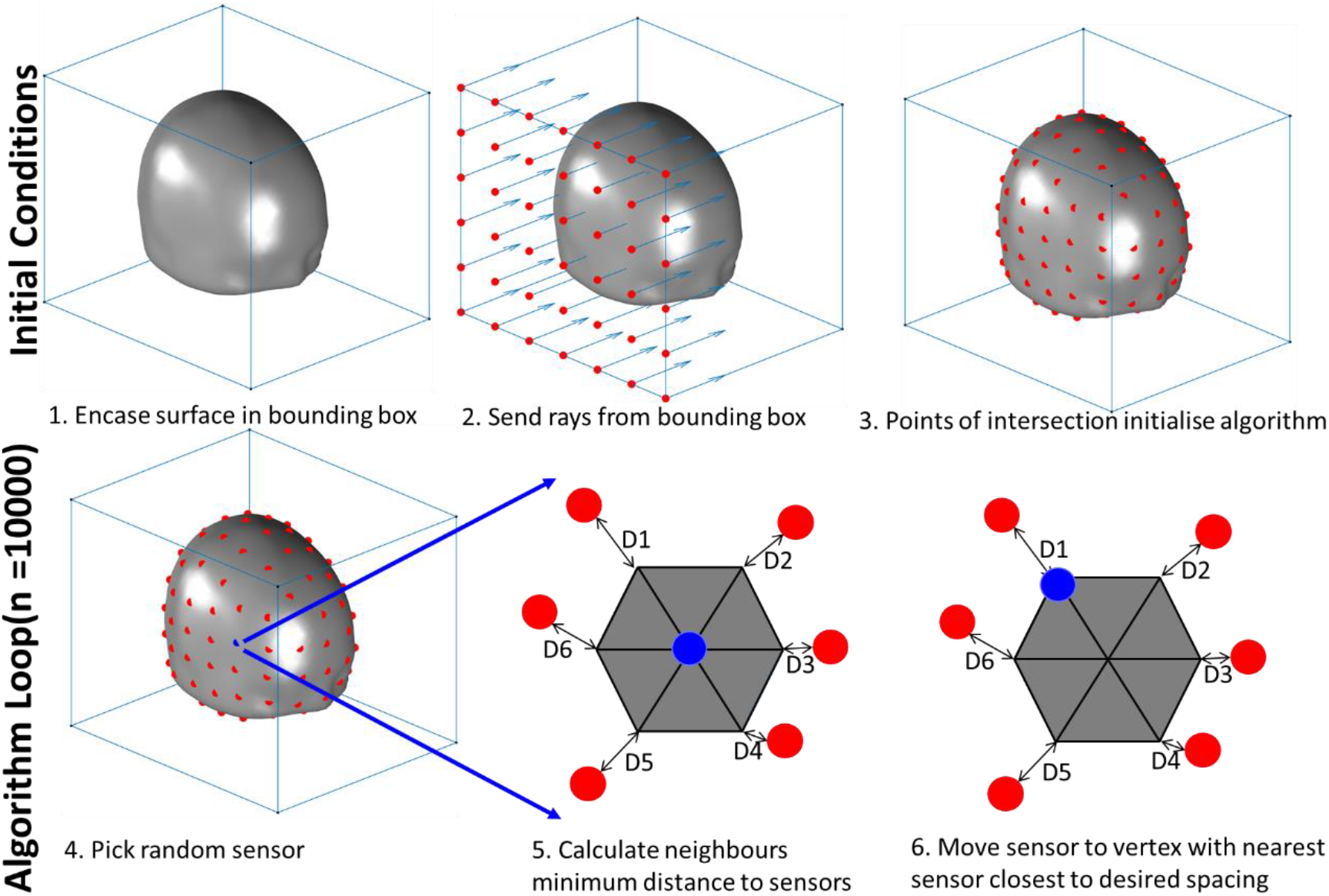
Algorithm for sensor placement. Initially the bounding box of the surface is subdivided into squares of edge length equal to the desired sampling density (panel 2). These points are then projected onto the surface to initialise the algorithm (panel 3). In the optimisation stage, at each loop iteration, a sensor is chosen at random (panel 4) and moved to a neighbouring vertex if the movement brings the observed sampling density closer to the target sampling denisty (panels 5, 6). This process is then repeated.

#### 2.3.1 The effect of SNR, depth and spatial sampling on spatial discrimination

We create 50 different arrays ranging in sampling density from 10mm to 60mm sampling density. The code required to create these arrays is available via GitHub (https://github.com/tierneytim/OPM).

We simulate MEG data as a sine wave with 10 Hz frequency. The SNR of the response was manipulated by changing the source amplitude from 1nAm to 10nAm to 100nAm assuming a 100ft noise standard deviation (uncorrelated across sensors) in all cases (corresponding to a 10 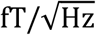 noise floor in 100 Hz bandwidth). The cortical surface normal is used to constrain the orientation of the source. The forward model was the Nolte single shell model (Nolte, 2003). There were 40 sources included in the simulation. The locations of the simulated sources were chosen to vary in depth and location across the whole brain. To meet this aim we selected the 20 deepest sources and the 20 most superficial sources in order to study the interaction, of depth, sampling density and SNR. To ensure the sources varied in position around the brain we ensured that no more than 1 source from each AAL region (Tzourio-Mazoyer et al., 2002) was chosen. The method of source inversion was an Empirical Bayesian Beamformer (Belardinelli, Ortiz, Barnes, Noppeney, & Preissl, 2012) as implemented in SPM12 (https://www.fil.ion.ucl.ac.uk/spm/). This method has been used in a similar context with real data to distinguish between models of cortical anatomy (López, Valencia, Flandin, Penny, & Barnes, 2017) and models with missing or displaced hippocampal anatomy (Tzovara et al., 2019). In order to relate our discrimination metrics to conventional cryogenic systems we used the same metrics to look at sensor arrays displaced 20mm from the scalp surface.

#### 2.3.2 The effect of gain changes on spatial discrimination

The gain errors were assumed to unrelated for each sensor and drawn from a normal distribution of mean 0 and standard deviation 1%, 2.5% and 10% of the nominal gain. We examine this effect across the three SNR levels which are set by fixing the source level currents (1nAm, 10nAm, and 100nAm) over the 40 brain regions varying in depth and location.

## 3. Results

### 3.1 Depth, SNR, spatial sampling and spatial discrimination: on scalp and off scalp

The results of simulations described in section **2.3.1** are graphically represented in Figure 3. The black contour lines indicate the point at which nearby sources can be discriminated for each candidate array. For example, for shallow sources (left column) at moderate SNR (middle row) the ability to discriminate sources at less than 5mm, reading from the F=3 (or p<0.05) contour line) would require> 200 channels (spatial sampling 20mm). However, if one wished to discriminate sources at 10mm (for the same moderate SNR at approximately 30mm spacing) only 87 sensors would be required. The same curves for typical cryogenic configurations (sensors offset from the scalp by 20mm) are shown in Supplementary figure 1. In this case the trends are the same except that more channels are required in order to reach the same discrimination performance.

**Figure 3.**
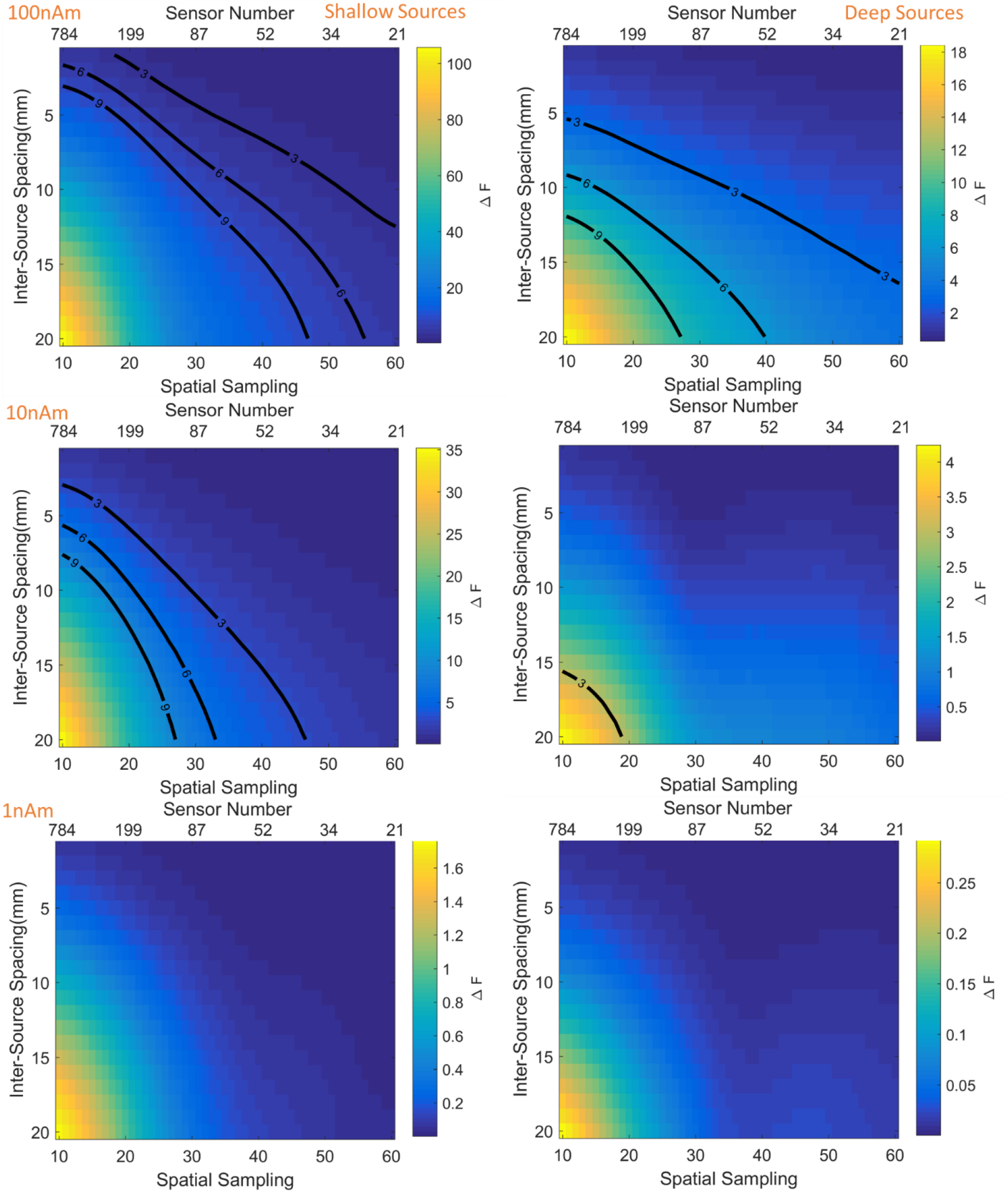
Discrimination between models as a function of spatial sampling density (or channel number). Left and right columns depict shallow and deep sources respectively. Rows show three different source amplitudes (100nAm, 10nAm, 1nAm). The colour scale shows the change in free energy relative to the base model. Thick black lines delineates the sensor spatial sampling necessary to confidently (p<0.05 or F>3) discriminate sources at a given inter source spacing.

### 3.2 Number of sensors

We can graph the relationship between spatial discrimination and spatial sampling for on and off scalp systems (shown here displaced by 20mm) at a given SNR (Figure 4). Regardless of the SNR it is found that to achieve the same spatial discrimination on scalp systems require fewer sensors than their off scalp counterparts. What is of interest is the disparity between how many sensors are needed in an on and off scalp system to achieve the same spatial discrimination. For example, to achieve 10 mm spatial discrimination between sources ~40 rather than ~70 channels are required for on-scalp vs. off - scalp sensors respectively at high SNR. While ~150 on scalp sensors are required for moderate SNR in comparison to > 500 sensors for the off scalp system. It is also clear that there are diminishing returns for having much more than 100 sensors in the on-scalp system as there is only a modest increase in spatial discrimination for each additional sensor.

**Figure 4.**
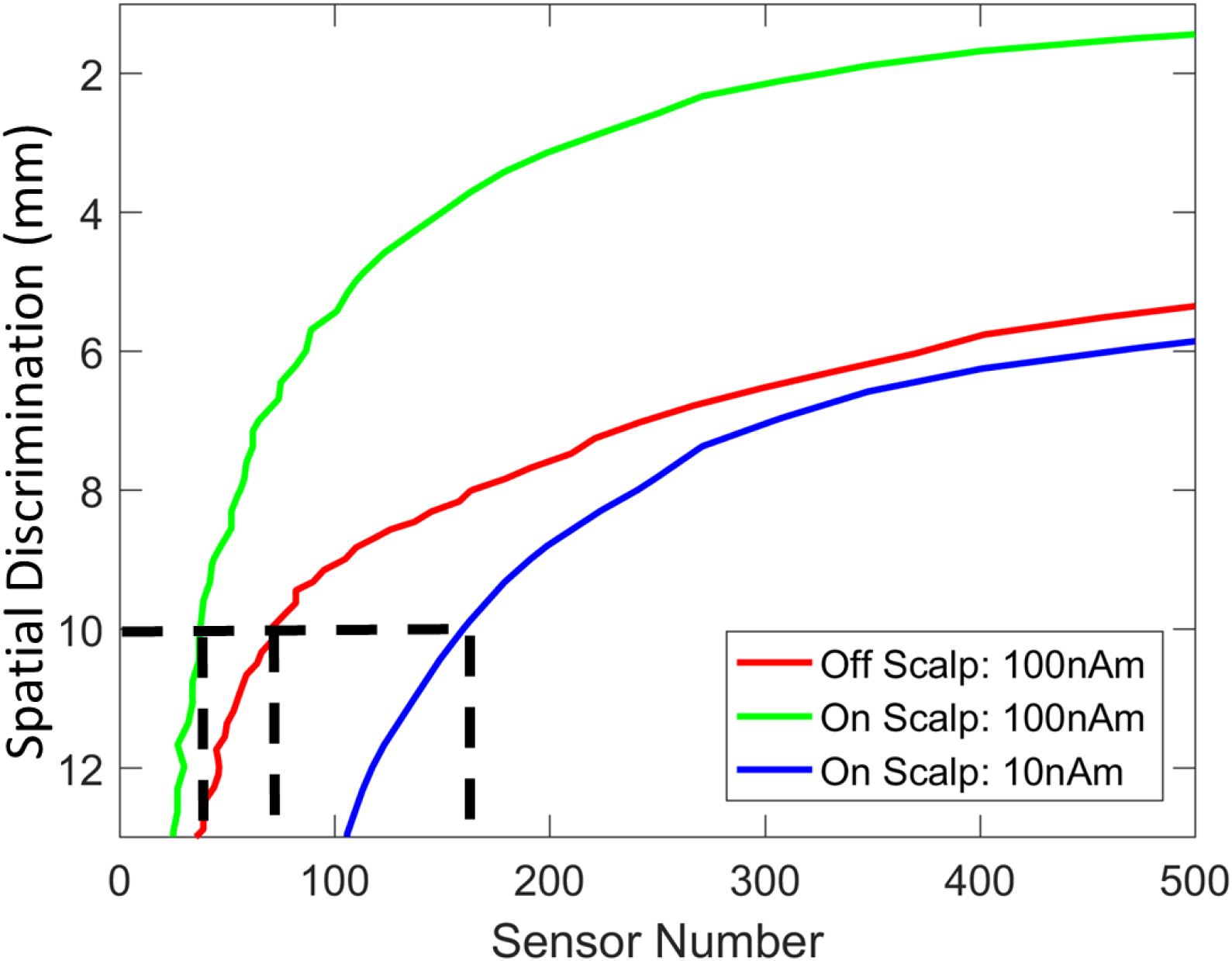
Sensor number and spatial discrimination for on- and off-scalp systems. Sensor number (x-axis) is graphed against spatial discrimination (y-axis) for both on-scalp (red, green) and off-scalp (blue) systems at different signal levels (100 nAm, 10nAm). Neither system had discrimination performance below 12mm at 1nAm and the off-scalp system did not reach this level at 10nAm and so these curves are not shown. While both systems are capable of achieving spatial discrimination of sources better than 1cm across the whole brain (black dotted lines), the on-scalp system (red and green curves) can achieve this with considerably fewer channels. Importantly, one can see for a given spatial discrimination an on-scalp system requires fewer sensors than an off-scalp system.

### 3.3: The impact of gain errors on spatial discrimination

From the previous section we can conclude that there are diminishing returns for having >100 on scalp sensors. As such for examining gain errors we use an on-scalp system with approximately 100 sensors corresponding to an approximate 30mm spacing. The simulations include 30 repetitions of 3 different gain levels (1%, 2.5% and 10%) for 3 different Signal levels (1nA, 10nA, 100nA with 100fT standard deviation sensor level noise). The results for deep and superficial sources are presented in Figure 5. It can be seen that the gain errors have the biggest impact at high SNR. At moderate and low levels of SNR the effects of gain errors are masked by the noise. Interestingly the gain errors appear more detrimental for deeper sources. Ultimately, though SNR is the dominant determinant of spatial discrimination with changes in spatial discrimination being only noticeable for very large gain errors (10%). These gain errors would be analogous to random sensor orientation errors of ~8, 13, and 26 degrees respectively. Similar relationships between gain errors, positional and orientation errors for smaller OPM systems are documented elsewhere (Duque-Munoz et al., 2019).

**Figure 5.**
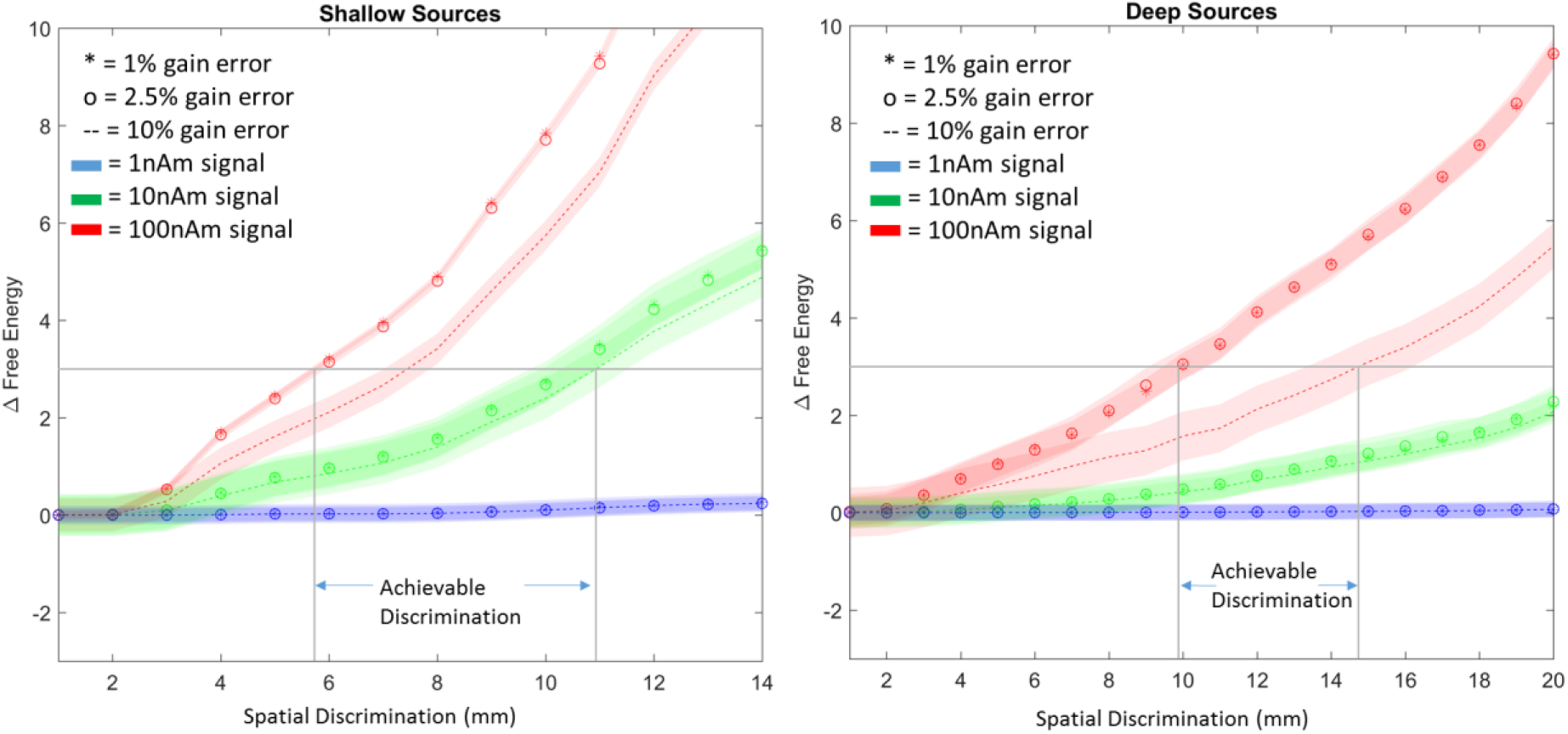
The effect of gain errors on spatial discrimination of OP-MEG systems. In the left panel results for shallow sources are presented. High, moderate and low SNR are signified by red green and blue, respectively. The thickness represents standard error. The x axis is spatial discrimination and the y axis is change in free energy. Stars represent 1% gain error, circles represent 2.5% gain error and dotted lines represent 10% gain error. The horizontal light grey lines represent the free energy threshold of 3 while the vertical grey lines show what the spatial discrimination is limited to in the presence of gain errors. The same symbols and colour schemes are used for the right panel to describe deep sources.

## 4. Discussion

In this paper we have presented a model comparison framework to quantify spatial discrimination of competing sensor arrays. We show that spatial discrimination of 1cm requires relatively few on-scalp sensors (<100) for high SNR for both deep and superficial sources. This corresponds to an inter-sensor distance of approximately 3cm. For this sensor layout, gain errors have the greatest impact on deep source discrimination at high SNR.

Our findings are in line with previous OPM simulation studies (Boto, Bowtell, et al., 2016). The main contribution of this study is the change to a probabilistic formalism in which the sensor array is dependent on the neuroscientific question/desired spatial discrimination. We should note that we are far from the theoretical limits of spatial discrimination for these on-scalp sensors. Indeed recent studies (M. S. Hämäläinen & Lundqvist, 2019; Iivanainen, Mäkinen, Zetter, et al., 2019) has shown that, due to the higher spatial frequency fields now measureable, to fully exploit the potential of OPM systems one would require of the order 300-500 sensors.

There are other ways to define spatial discrimination such as low correlation between sources (Boto, Bowtell, et al., 2016; Iivanainen et al., 2017) or examination of point-spread functions considered resolved. As such, defining spatial discrimination is arguably a statistical problem. The proposed framework has also been used extensively to model laminar sources (Bonaiuto et al., 2017, 2018; Troebinger et al., 2014) in MEG as well as make probabilistic statements about deep structure activity (Khemka, Barnes, Dolan, & Bach, 2017; Meyer et al., 2017b) where spatial precision is paramount.

SNR is an important factor in these simulations and it becomes clear that lower sensor numbers can often answer the same question if the source strength is high enough. As we can achieve higher SNR by simply recording for longer and averaging over more data (Brookes et al., 2010), fewer sensors can be traded against longer recording times. Indeed most OPM experiments to date (that attempt a source reconstruction) have been performed with far fewer (typical 16-32) sensors than a traditional cryogenic system (Barry et al., 2019; Boto et al., 2018; Boto, Meyer, et al., 2016; Holmes et al., 2018; Iivanainen, Zetter, & Parkkonen, 2019; Lin et al., 2019; Roberts et al., 2019; Tierney et al., 2018; Zetter, Iivanainen, & Parkkonen, 2019).

We have made several idealistic assumptions. Practically, achievable discrimination performance may be bounded by other factors. For example it has been shown that forward modelling errors limit discrimination at high SNR (Boto, Bowtell, et al., 2016; Arjan Hillebrand & Barnes, 2003). Here, we assume that the source model is known and we have created and reconstructed the data under the same forward models. Other studies have looked at these models in more detail (Stenroos, Hunold, & Haueisen, 2014; Stenroos & Sarvas, 2012) and we do not examine this factor directly. That said, increasing the channel number will not solve these issues. We had initially expected gain errors to have similar consequences to forward modelling errors but it appears these are much less pernicious. This is because the gain changes have been considered independent across sensors (rather than systematic for lead-field errors) and indeed we found that for practical OPM gain errors (+−2%) there were negligible effects on discrimination performance. However, there may be instances when gain changes have some correlated spatial structure across the array (for example as the subject moves through a magnetic field gradient) and this would most likely be more detrimental to the source modelling.

We have also made the assumption that our model of the cortical current flow (and its covariance) is known. The ideal inversion algorithm for a particular question or disease state (like epilepsy) is far from decided, however the model evidence framework outlined here could also be used to optimize inversion assumptions (Friston et al., 2008) as well as measurement arrays.

We have provided a simulation based framework for users of OPMs to establish how many sensors are required for neuroscientific applications to achieve a desired spatial discrimination. We find adequate discrimination (<1cm) can be easily obtained with few sensors (<100) at coarse spatial samplings (>30mm) at high SNR for both deep and superficial sources. We also demonstrate that the dominant determinant of spatial discrimination in an OPM system is not gain changes/ calibration errors but the magnitude of the physiological response and the depth of the source.

