## Supplemental figure 1 for "Pragmatic spatial sampling for wearable MEG arrays"

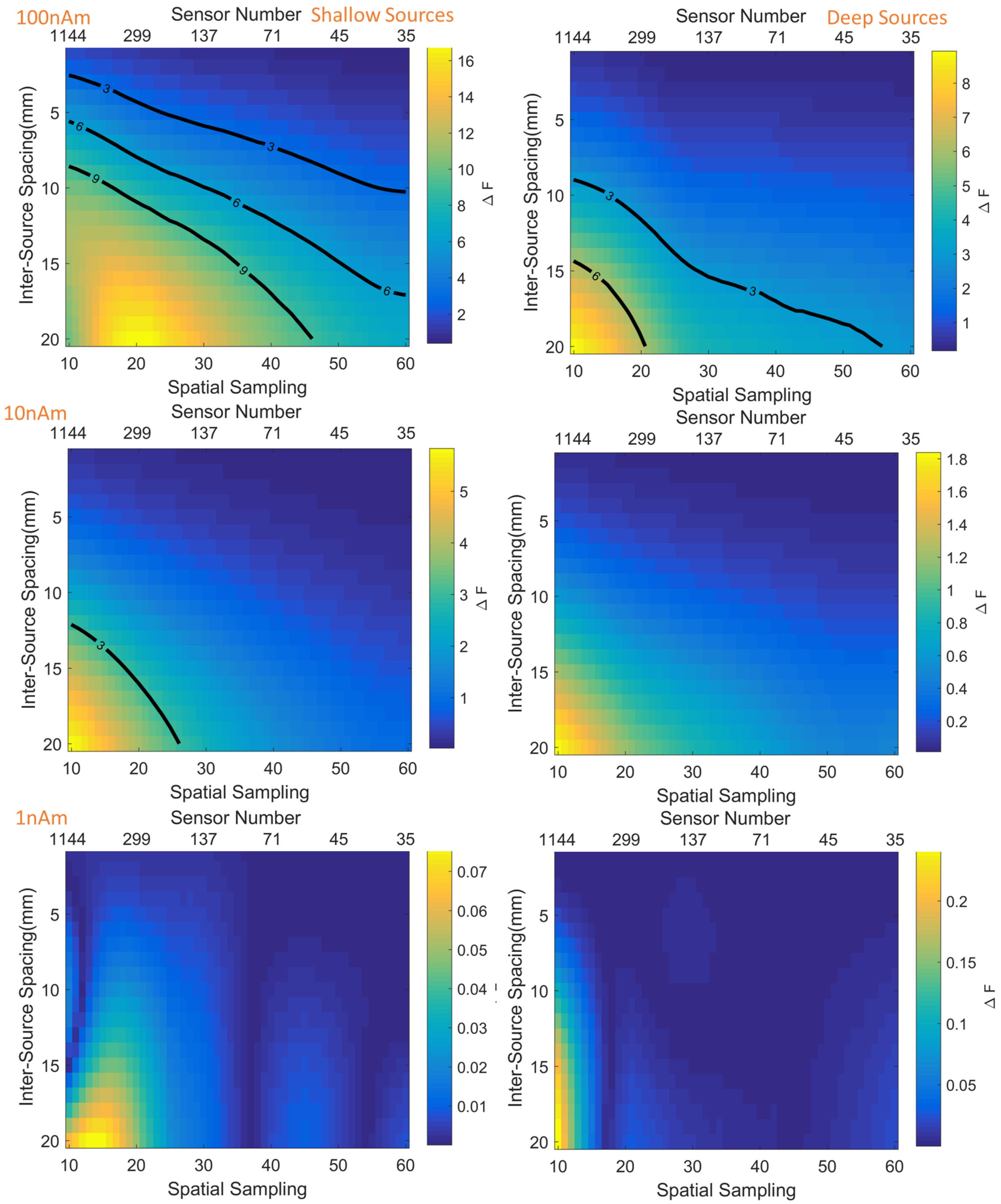


Figure S1. Discrimination between models as a function of spatial sampling density (or channel number) for off scalp systems (20mm). Left and right columns depict shallow and deep sources respectively. Rows show three different source amplitudes (100nAm, 10nAm, 1nAm). The colour scale shows the change in free energy relative to the base model. Thick black lines delineates the spatial sampling necessary to confidently ($p<0.05$) discriminate sources at a given inter source spacing.
